# Synthesis and Biological Evaluation of Novel MB327 Analogs as Resensitizers for Desensitized Nicotinic Acetylcholine Receptors after Intoxication with Nerve Agents

**DOI:** 10.1101/2024.02.09.579646

**Authors:** Tamara Bernauer, Valentin Nitsche, Jesko Kaiser, Christoph G.W. Gertzen, Georg Höfner, Karin V. Niessen, Thomas Seeger, Dirk Steinritz, Franz Worek, Holger Gohlke, Klaus T. Wanner, Franz F. Paintner

## Abstract

Poisoning with organophosphorus compounds, which can lead to a cholinergic crisis due to the inhibition of acetylcholinesterase and the subsequent accumulation of acetylcholine (ACh) in the synaptic cleft, is a serious problem for which treatment options are currently insufficient. Our approach to broadening the therapeutic spectrum is to use agents that interact directly with desensitized nicotinic acetylcholine receptors (nAChRs) in order to induce functional recovery after ACh overstimulation. Although MB327, one of the most prominent compounds investigated in this context, has already shown positive properties in terms of muscle force recovery, this compound is not suitable for use as a therapeutic agent due to its insufficient potency. By means of *in silico* studies based on our recently presented allosteric binding pocket at the nAChR, i.e. the MB327-PAM-1 binding site, three promising 4-aminopyridinium ion-substituted MB327 analogs (PTM0056, PTM0062 and PTM0063) were identified. In this study, we present the synthesis and biological evaluation of a series of new 4-aminopyridinium ion-substituted analogs of the aforementioned compounds (PTM0064-PTM0072), as well as hydroxy-substituted analogs of MB327 (PTMD90-0012 and PTMD90-0015) designed to substitute energetically unfavorable water clusters identified during molecular dynamics simulations. The compounds were characterized in terms of their binding affinity towards the aforementioned binding site by applying the UNC0642 MS Binding Assays and in terms of their muscle force reactivation in rat diaphragm myography. More potent compounds were identified compared to MB327, as some of them showed a higher affinity towards MB327-PAM-1 and also a higher recovery of neuromuscular transmission at lower compound concentrations. To improve the treatment of organophosphate poisoning, direct targeting of nAChRs with appropriate compounds is a key step, and this study is an important contribution to this research.

## 1 Introduction

The verified use of sarin in the Syrian war in 2013 (Dolgin, 2013; Pita and Domingo, 2014) and 2017 (OPCW, 2017), as well as the politically motivated Novichok attack on Russian opposition politician Alexei Navalny in 2020 (Steindl et al., 2021), provide clear evidence that highly toxic organophosphorus nerve agents still pose a major threat to military personnel and civilians, even after their international ban by the Chemical Weapons Convention in 1997 (Thakur and Haru, 2007). Intoxication with organophosphorus compounds (OPCs) leads to the irreversible inhibition of acetylcholinesterase (AChE), resulting in an uncontrolled accumulation of acetylcholine (ACh) in the synaptic cleft of cholinergic neurons. As a result, cholinergic signaling is disrupted by overstimulation of muscarinic and nicotinic acetylcholine receptors (mAChRs and nAChRs, respectively) (Koelle, 1981; Maselli and Leung, 1993; Massoulié et al., 1993). This condition, known as a “cholinergic crisis”, can be life-threatening due to respiratory paralysis as a result of the disruption of nAChR-mediated neuromuscular transmission (Brown and Brix, 1998; Thiermann et al., 2010). Standard treatment for nerve agent poisoning currently includes a muscarinic acetylcholine receptor antagonist, e.g. atropine, to reduce mAChR overstimulation and an oxime-based AChE reactivator, e.g. obidoxime. According to Sheridan et al. (Sheridan et al., 2005) neuromuscular blockers (competitive nAChR antagonists) are not suitable for the treatment of nerve agent poisoning. Hence, reactivation of inhibited AChE appears to be crucial to counteract nicotinic overstimulation. However, despite decades of effort, there is still no universally applicable AChE reactivator that can efficiently cleave all OPC-AChE conjugates (Worek et al., 2020). Therefore, there is an urgent need to develop novel antidotes to counteract desensitization of muscle-type nAChRs as a result of overstimulation. One approach is to use agents that directly target the nAChR to restore its function after it has been desensitized by overstimulation (Sheridan et al., 2005; Turner et al., 2011).

Indeed, a number of bispyridinium salts, such as the prototypical compound MB327 (Figure 1), are able to restore the function of desensitized nAChR of *Torpedo californica* (recently reclassified as *Tetronarce californica*) in *in vitro* experiments by interacting directly with the receptor most likely via an allosteric mechanism (Niessen et al., 2016; Niessen et al., 2018; Seeger et al., 2012; Sichler et al., 2018). Since the *Torpedo californica* nAChR has a high sequence identity to the rat and human muscle-type nAChRs, it is reasonable to assume that MB327 has similar effects on the desensitized nAChRs of the latter species as well. Indeed, in *ex vivo* experiments, MB327 shows a muscle force-restoring effect on both soman-poisoned rat diaphragms and soman-poisoned human intercostal muscles, which is thought to be due to the resensitization of desensitized nAChR (Niessen et al., 2018; Seeger et al., 2012). In addition, MB327 (or the corresponding methanesulfonate salt MB399, respectively) in combination with the mAChR antagonist hyoscine and the indirect parasympathomimetic physostigmine, was shown to increase the survival rates of guinea pigs poisoned with sarin or tabun in *in vivo* studies (Timperley et al., 2012; Turner et al., 2011). Despite these promising results, MB327 is not suitable for human use due to its narrow therapeutic window. Nevertheless, MB327 is a promising starting point for further investigation.

**Figure 1.**
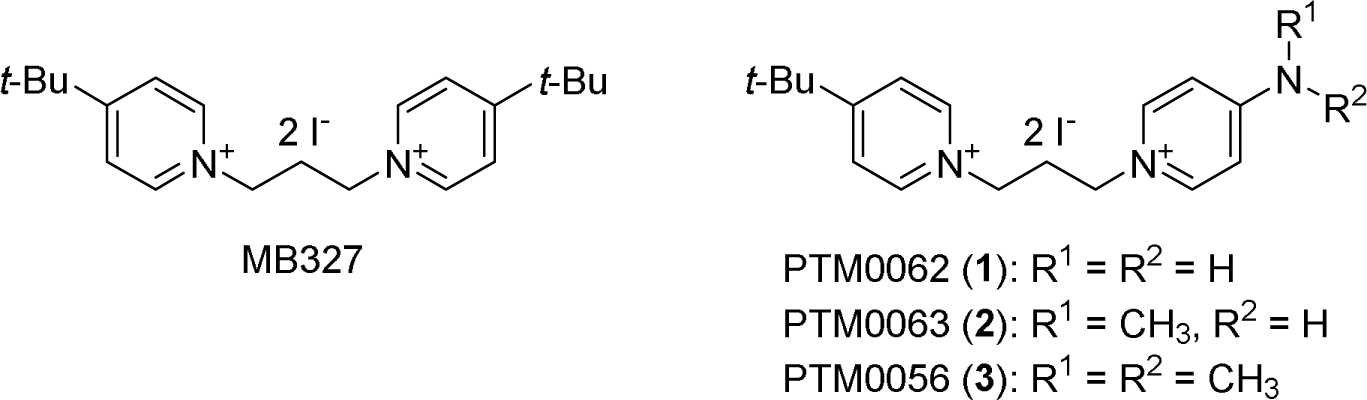
Chemical structures: MB327, PTM0062 (**1**), PTM0063 (**2**) and PTM0056 (**3**).

Recently, based on blind docking experiments and molecular dynamics simulations, we proposed a new allosteric binding site of MB327 at the muscle-type nAChR, termed MB327-PAM-1 (Kaiser et al., 2023). This binding site is located at the transition from the extracellular to the trans-membrane region and, according to a rigidity analysis, is expected to exert an allosteric effect on the orthosteric binding pocket upon binding of MB327. The amino acids interacting with MB327 in this binding site, predominantly glutamate residues, are highly conserved within the different nAChR subunits as well as in different species. Accordingly, comparable resensitizing effects of MB327 should be observable on desensitized nAChR of different species (e.g. *Torpedo californica*, rats and humans).

However, recently published results also show that bispyridinium compounds related to MB327 have inhibitory activity on the nAChR, most likely mediated via the orthosteric binding site (Epstein et al., 2021). Indeed, free ligand diffusion MD simulations performed by us indicated that MB327 also has affinity for the orthosteric binding site (Kaiser et al., 2023). The fact that the muscle force restoring activity of MB327 on soman-poisoned rat diaphragms is lost at higher concentrations after a peak at 300 µM may be explained by this inhibitory activity (Niessen et al., 2018).

The binding mode of MB327 in the MB327-PAM-1 binding pocket indicates that one of the two *tert*-butyl groups projects into a polar region of the binding pocket. This allowed us to predict structural modifications of MB327 that led to the more potent resensitizers PTM0062 (**1**) and PTM0063 (**2**), which have a more polar substituent, i.e., an amino and a methylamino group, respectively, instead of one of the two *tert*-butyl residues of MB327 (Figure 1) (Kaiser et al., 2023). Interestingly, the recently described dimethylamino analog PTM0056 (**3**) (Figure 1), which has a less polar substituent than PTM0062 (**1**) or PTM0063 (**2**), also shows a slightly higher affinity for the MB327-PAM-1 binding site than MB327 (Rappenglück et al., 2018) and a muscle force-restoring activity comparable to PTM0062 (**1**) and PTM0063 (**2**) in preliminary, unpublished *ex vivo* studies with soman-poisoned rat diaphragm hemispheres. Therefore, PTM0056 (**3**) appeared to be a promising starting point for the development of new, possibly even more potent, resensitizers for the desensitizied muscle-typ nAChR.

In this study, we report the development of a series of non-symmetric MB327 analogs derived from PTM0062 (**1**), PTM0063 (**2**) and PTM0056 (**3**), respectively, as resensitizers for desensitized muscle-type nAChRs. To obtain a structurally diverse set of analogs, the 4-amino substituents of compounds **1**-**3** were replaced by either acylamino groups, new dialkylamino groups or cyclic amino groups (e.g. pyrrolidino, piperidino, morpholino and piperazino groups). Furthermore, based on identifying energetically unfavorable water clusters in MB327-PAM-1 using molecular dynamics (MD) simulations in combination with Grid Inhomogeneous Solvent Theory (GIST) computations (Lazaridis, 1998; Nguyen et al., 2011; Nguyen et al., 2012; Ramsey et al., 2016), we designed novel MB327 analogs potentially able to substitute these water clusters. To determine the affinities of the newly developed compounds at the MB327-PAM-1 binding site of *Torpedo*-nAChR, our recently developed UNC0642 MS Binding Assays were applied. To gain insight into the intrinsic activity of some selected representatives of the newly developed compounds, their ability to restore muscle force was also investigated in *ex vivo* experiments with soman-poisoned rat diaphragms. Some of the newly developed compounds showed a slightly higher affinity for the MB327-PAM-1 binding site of *Torpedo*-nAChR than the prototypical compound MB327 and a comparable or even higher muscle force-restoring effect on soman-poisoned rat diaphragms at lower concentrations than MB327.

## 2 Material and Methods

### 2.1 Synthesis of novel MB327 analogs

Microwave reactions were carried out on a Discover SP microwave system by *CEM GmbH* in glass vials. All chemicals were used as purchased from commercial sources. Solvents used for crystallization were distilled before use. Melting points were determined with a Büchi 510 melting point apparatus and are uncorrected. For IR spectroscopy, an FT-IR Spectrometer 1600 from *PerkinElmer* was used. High-resolution mass spectrometry was performed on a Finnigan MAT 95 (EI) or a Finnigan LTQ FT (ESI). ^1^H and ^13^C NMR spectra were recorded on a Bruker BioSpin Avance III HD 400 and 500 MHz at 25 °C. For data processing, MestReNova (Version 14.1.0) from Mestrelab Research S.L. 2019, and for calibration, the solvent signal (CD_3_OD) was used. Using PTM0064 (**6a**) as an example, the signal assignment in the NMR spectra of the bispyridinium compounds is shown (see Supporting information). Unless otherwise noted, the purity of the test compounds was ≥ 98%, determined using quantitative ^1^H NMR spectroscopy using TraceCERTS® ethyl 4-(dimethylamino)benzoate from *Sigma Aldrich* as internal calibrant (Cushman et al., 2014; Pauli et al., 2014).

All target compounds synthesized in the context of this study were cataloged with a certain PTM and PTMD number, respectively (<u>P</u>harmacy and <u>T</u>oxicology <u>M</u>unich and <u>P</u>harmacy and <u>T</u>oxicology <u>M</u>unich and <u>D</u>üsseldorf, respectively).

#### General Procedure (GP): Synthesis of non-symmetric MB327 analogs by *N*-alkylation with 4

A solution of 4-(*tert*-butyl)-1-(3-iodopropyl)pyridin-1-ium iodide (**4**) (Rappenglück et al., 2018) (1.0 equiv) and the corresponding 4-amino-substituted pyridine **5** (1.05-1.1 equiv) or 7-hydroxychinoline (**7**) in acetonitrile (2.0-2.7 mL/mmol) was stirred at 90 °C under microwave irradiation (150 W) for 1 h unless otherwise stated. The reaction mixture was concentrated in vacuo, and the residue was purified by recrystallization from different solvent mixtures.

**4-Acetamido-1-{3-[4-(*tert*-butyl)pyridin-1-ium-1-yl]propyl}pyridin-1-ium diiodide (PTM0064, 6a):** According to the GP, with **4** (431 mg, 1.00 mmol, 1.0 equiv) and **5a** (150 mg, 1.10 mmol, 1.1 equiv) in MeCN (2.0 mL). Recrystallization from EtOAc/EtOH/MeOH (1:2:1) afforded **6a** (529 mg, 93%) as a yellow solid. m.p. 259 °C; ^1^H NMR (400 MHz, CD_3_OD): δ = 1.45 (s, 9 H), 2.27 (s, 3 H), 2.71-2.80 (m, 2 H), 4.66-4.72 (m, 2 H), 4.77-4.83 (m, 2 H), 8.12-8.16 (m, 2 H), 8.16-8.20 (m, 2 H), 8.79-8.86 (m, 2 H), 8.96-9.02 (m, 2 H); ^13^C NMR (101 MHz, CD_3_OD): δ = 24.61, 30.21, 33.27, 37.68, 57.42, 58.32, 116.48, 126.90, 145.43, 146.32, 154.33, 172.69, 173.27; IR (KBr): L = 2964, 1645, 1518, 1207 cm^-1^; HRMS (ESI): *m/z* calcd for C_19_H_27_N_3_OI_2_-I^-^: 440.1199 [*M*-I]^+^; found: 440.1185.

**4-[(*tert*-Butoxycarbonyl)amino]-1-{3-[4-(*tert*-butyl)pyridin-1-ium-1-yl]propyl}-pyridin-1-ium diiodide (PTM0065, 6b):** According to the GP, a solution of **4** (323 mg, 0.750 mmol, 1.0 equiv) and **5b** (157 mg, 0.810 mmol, 1.08 equiv) in MeCN (2.0 mL) was stirred under microwave irradiation (150 W) at 60 °C for 15 h. Recrystallization from Et_2_O/DMF (10:1) afforded **6b** (431 mg, 92%) as a yellow solid. Purity: 94%. m.p. 117 °C; ^1^H NMR (400 MHz, CD_3_OD): δ = 1.45 (s, 9 H), 1.56 (s, 9 H), 2.67-2.79 (m, 2 H), 4.59-4.67 (m, 2 H), 4.74-4.82 (m, 2 H), 7.87-8.02 (m, 2 H), 8.20-8.14 (m, 2 H), 8.68-8.76 (m, 2 H), 8.91-9.01 (m, 2 H); ^13^C NMR (101 MHz, CD_3_OD): δ = 28.25, 30.21, 33.19, 37.68, 57.16, 58.38, 84.27, 115.30, 126.90, 145.41, 145.85, 153.10, 155.64, 173.30; IR (KBr): L = 2967, 1643, 1531, 1148 cm^-1^; HRMS (ESI): *m/z* calcd for C_22_H_33_N_3_O_2_I_2_-I^-^: 498.1618 [*M*-I]^+^; found: 498.1600.

**4-[Benzyl(methyl)amino]-1-{3-[4-(*tert*-butyl)pyridin-1-ium-1-yl]propyl}pyridin-1-ium diiodide (PTM0066, 6c):** According to the GP, with **4** (323 mg, 0.750 mmol, 1.0 equiv) and **5c** (164 mg, 0.830 mmol, 1.1 equiv) in MeCN (2.0 mL). Recrystallization from EtOAc/EtOH/MeOH (10:5:1) afforded **6c** (354 mg, 75%) as a yellow solid. m.p. 185 °C; ^1^H NMR (400 MHz, CD_3_OD): δ = 1.45 (s, 9 H), 2.56-2.70 (m, 2 H), 3.34 (s, 3 H), 4.40 (t, *J* = 7.6 Hz, 1 H), 4.74 (t, *J* = 7.7 Hz, 2 H), 4.90 (s, 2 H), 7.03-7.19 (m, 2 H), 7.24-7.29 (m, 2 H), 7.29-7.41 (m, 3 H), 8.13-8.19 (m, 2 H), 8.20-8.40 (m, 2 H), 8.91-8.97 (m, 2 H); ^13^C NMR (101 MHz, CD_3_OD): δ = 30.20, 33.14, 37.67, 39.70, 55.41, 56.75, 58.52, 109.83, 126.86, 127.82, 129.11, 130.21, 136.21, 143.72, 145.38, 158.26, 173.24; IR (KBr): L = 2967, 1645, 1556, 1193 cm^-1^; HRMS (ESI): *m/z* calcd for C_25_H_33_N_3_I_2_-I^-^: 502.1719 [*M*-I]^+^; found: 502.1703.

**4-(*tert*-Butyl)-1-{3-[4-(diethylamino)pyridin-1-ium-1-yl]propyl}pyridin-1-ium diiodide (PTM0067, 6d):** According to the GP, with **4** (431 mg, 1.00 mmol, 1.0 equiv) and **5d** (165 mg, 1.10 mmol, 1.1 equiv) in MeCN (2.0 mL). Recrystallization from EtOAc/EtOH (2.5:1) afforded **6d** (419 mg, 72%) as a yellow solid. m.p. 217 °C; ^1^H NMR (400 MHz, CD_3_OD): δ = 1.27 (t, *J* = 7.2 Hz, 6 H), 1.45 (s, 9 H), 2.60-2.71 (m, 2 H), 3.64 (q, *J* = 7.2 Hz, 4 H), 4.41 (t, *J* = 7.7 Hz, 2 H), 4.78 (t, *J* = 7.8 Hz, 2 H), 7.01-7.08 (m, 2 H), 8.13-8.21 (m, 2 H), 8.24-8.31 (m, 2 H), 8.97-9.02 (m, 2 H); ^13^C NMR (101 MHz, CD_3_OD): δ = 12.21, 30.23, 33.20, 37.66, 46.46, 55.15, 58.49, 109.23, 126.86, 143.44, 145.41, 156.42, 173.11; IR (KBr): L = 2968, 1648, 1561, 1197 cm^-1^; HRMS (ESI): *m/z* calcd for C_21_H_33_N_3_I_2_-I^-^: 454.1719 [*M*-I]^+^; found: 454.1704.

**4-(*tert*-Butyl)-1-{3-[4-(pyrrolidin-1-yl)pyridin-1-ium-1-yl]propyl}pyridin-1-ium diiodide (PTM0068, 6e):** According to the GP, with **4** (431 mg, 1.00 mmol, 1.0 equiv) and **5e** (159 mg, 1.05 mmol, 1.05 equiv) in MeCN (2.0 mL). Recrystallization from EtOAc/*i*-PrOH (1.1:1) afforded **6e** (457 mg, 79%) as a yellow solid. m.p. 178 °C; ^1^H NMR (400 MHz, CD_3_OD): δ = 1.45 (s, 9 H), 2.10-2.15 (m, 4 H), 2.60-2.69 (m, 2 H), 3.55-3.60 (m, 4 H), 4.40 (t, *J* = 7.7 Hz, 2 H), 4.76 (t, *J* = 7.8 Hz, 2 H), 6.86-6.91 (m, 2 H), 8.15-8.18 (m, 2 H), 8.24-8.29 (m, 2 H), 8.96-9.00 (m, 2 H); ^13^C NMR (101 MHz, CD_3_OD): δ = 26.14, 30.21, 33.22, 37.66, 49.83, 55.26, 58.52, 109.87, 126.86, 143.01, 145.40, 155.23, 173.15; IR (KBr): L = 2961, 1647, 1561, 1192 cm^-1^; HRMS (ESI): *m/z* calcd for C_21_H_31_N_3_I_2_-I^-^: 452.1563 [*M*-I]^+^; found: 452.1545.

**4-(*tert*-Butyl)-1-{3-[4-(piperidin-1-yl)pyridin-1-ium-1-yl]propyl}pyridin-1-ium diiodide (PTM0069, 6f):** According to the GP, with **4** (216 mg, 0.500 mmol, 1.0 equiv) and **5f** (85.2 mg, 0.530 mmol, 1.05 equiv) in MeCN (1.0 mL). Recrystallization from EtOAc/*i*-PrOH (1:1.2) afforded **6f** (245 mg, 83%) as a yellow solid. m.p. 240 °C; ^1^H NMR (400 MHz, CD_3_OD): δ = 1.45 (s, 9 H), 1.69-1.76 (m, 4 H), 1.76-1.85 (m, 2 H), 2.58-2.71 (m, 2 H), 3.69-3.76 (m, 4 H), 4.39 (t, *J* = 7.6 Hz, 2 H), 4.77 (t, *J* = 7.7 Hz, 4 H), 7.12-7.20 (m, 2 H), 8.13-8.20 (m, 2 H), 8.21-8.29 (m, 2 H), 8.95-9.02 (m, 2 H); ^13^C NMR (101 MHz, CD_3_OD): δ = 24.93, 26.73, 30.22, 33.14, 37.66, 49.17, 55.07, 58.51, 109.45, 126.86, 143.58, 145.41, 156.92, 173.13; IR (KBr): L = 2956, 1648, 1547, 1194 cm^-1^; HRMS (ESI): *m/z* calcd for C_22_H_33_N_3_I_2_-I^-^: 466.1719 [*M*-I]^+^; found: 466.1711.

**4-(*tert*-Butyl)-1-[3-(4-morpholinopyridin-1-ium-1-yl)propyl]pyridin-1-ium diiodide (PTM0070, 6g):** According to the GP, with **4** (431 mg, 1.00 mmol, 1.00 equiv) and **5g** (172 mg, 1.10 mmol, 1.05 equiv) in MeCN (2.0 mL). Recrystallization from EtOAc/EtOH (1:1.4) afforded **6g** (518 mg, 87%) as a yellow solid. m.p. 222 °C; ^1^H NMR (400 MHz, CD_3_OD): δ = 1.45 (s, 9 H), 2.62-2.72 (m, 2 H), 3.69-3.76 (m, 4 H), 3.80-3.85 (m, 4 H), 4.45 (t, *J* = 7.7 Hz, 2 H), 4.78 (t, *J* = 7.8 Hz, 2 H), 7.13-7.27 (m, 2 H), 8.14-8.20 (m, 2 H), 8.33-8.40 (m, 2 H), 8.97-9.04 (m, 2 H); ^13^C NMR (101 MHz, CD_3_OD): δ = 30.23, 33.21, 37.66, 47.73, 55.33, 58.44, 67.15, 109.72, 126.86, 143.86, 145.41, 157.85, 173.11; IR (KBr): L = 2964, 1650, 1545, 1193 cm^-1^; HRMS (ESI): *m/z* calcd for C_21_H_31_N_3_OI_2_-I^-^: 468.1512 [*M*-I]^+^; found: 468.1493.

4-[4-(*tert*-Butoxycarbonyl)piperazin-1-yl]-1-{3-[4-(*tert*-butyl)pyridin-1-ium-1-yl]propyl}-pyridin-1-ium diiodide (PTM0071, 6h): According to the GP, with **4** (216 mg, 0.500 mmol, 1.0 equiv) and **5h** (145 mg, 0.550 mmol, 1.1 equiv) in MeCN (1.0 mL). Recrystallization from EtOAc/EtOH (2:1) afforded **6h** (320 mg, 92%) as a yellow solid. m.p. 149 °C; ^1^H NMR (500 MHz, CD_3_OD): δ = 1.45 (s, 9 H), 1.49 (s, 9 H), 2.61-2.69 (m, 2 H), 3.62-3.68 (m, 4 H), 3.76-3.80 (m, 4 H), 4.42 (t, *J* = 7.6 Hz, 2 H), 4.75 (t, *J* = 7.7 Hz, 2 H), 7.19-7.23 (m, 2 H), 8.15-8.19 (m, 2 H), 8.31-8.36 (m, 2 H), 8.95-8.99 (m, 2 H); ^13^C NMR (126 MHz, CD_3_OD): δ = 28.59, 30.21, 33.17, 37.67, 43.60, 47.12, 55.38, 58.48, 81.95, 109.85, 126.86, 143.79, 145.40, 156.17, 157.71, 173.22; IR (KBr): L = 2967, 1649, 1416, 1169 cm^-1^; HRMS (ESI): *m/z* calcd for C_26_H_40_N_4_O_2_I_2_-I^-^: 567.2196 [*M*-I]^+^; found: 567.2180.

**4-(1-{3-[4-(*tert*-Butyl)pyridin-1-ium-1-yl]propyl}pyridin-1-ium-4-yl)piperazin-1-ium triiodide (PTM0072, 6i):** A solution of **6h** (69.4 mg, 0.100 mmol, 1.0 equiv) and iodo(trimethyl)silane (80.0 mg, 0.400 mmol, 4.0 equiv) in MeCN (2.0 mL) was stirred under argon at rt for 2 h. **6i** (71.3 mg, 99%) was afforded after filtration of the reaction mixture as a yellow solid. m.p. 109 °C; ^1^H NMR (400 MHz, CD_3_OD): δ = 1.45 (s, 9 H), 2.65-2.75 (m, 2 H), 3.47-3.55 (m, 4 H), 4.03-4.12 (m, 4 H), 4.52 (t, *J* = 7.5 Hz, 2 H), 4.75-4.86 (m, 2 H), 7.33-7.39 (m, 2 H), 8.14-8.20 (m, 2 H), 8.45-8.52 (m, 2 H), 9.01-9.06 (m, 2 H); ^13^C NMR (101 MHz, CD_3_OD): δ = 30.23, 33.27, 37.65, 44.04, 44.65, 55.66, 58.33, 110.83, 126.84, 144.37, 145.44, 158.02, 173.07; IR (KBr): L = 2963, 1647, 1549, 1191 cm^-1^; HRMS (ESI): *m/z* calcd for C_21_H_33_N_4_I_3_-H^+^-2I^-^: 467.1671 [*M*-I-I-H]^+^; found: 467.1663.

**1-{3-[4-(*tert*-Butyl)pyridin-1-ium-1-yl]propyl}-7-hydroxyquinolin-1-ium diiodide (PTMD90-0012, 8):** According to the GP, a solution of **4** (431 mg, 1.00 mmol, 1.0 equiv) and **7** (163 mg, 1.10 mmol, 1.1 equiv) in MeCN (2.0 mL) was stirred under microwave irradiation (150 W) at 90 °C for 3 h. Recrystallization from EtOH/EtOAc (2:1.5) afforded **8** (167 mg, 29%) as a yellow solid. m.p. 263 °C; ^1^H NMR (400 MHz, CD_3_OD): δ = 1.45 (s, 9 H), 2.76-2.89 (m, 2 H), 4.89-4.96 (m, 2 H), 5.09-5.19 (m, 2 H), 7.56 (dd, *J* = 9.0, 2.1 Hz, 1 H), 7.68 (d, *J* = 2.2 Hz, 1 H), 7.81 (dd, *J* = 8.1, 6.1 Hz, 1 H), 8.11-8.22 (m, 2 H), 8.29 (d, *J* = 9.1 Hz, 1 H), 8.95-9.04 (m, 3 H), 9.31 (dd, *J* = 6.1, 1.4 Hz, 1 H); ^13^C NMR (101 MHz, CD_3_OD): δ = 30.20, 31.59, 37.68, 54.85, 58.50, 101.59, 119.39, 123.77, 126.81, 126.88, 134.57, 142.53, 145.41, 148.20, 148.99, 167.19, 173.30; IR (film): L = 2968, 1628, 1209, 849 cm^-1^; HRMS (ESI): *m/z* calcd for C_21_H_26_N_2_OI_2_-H^+^-2I^-^: 321.1967 [*M*-I-I-H]^+^; found: 321.1961.

1,1’-(2-Hydroxypropan-1,3-diyl)bis[4-(*tert*-butyl)pyridin-1-ium] dibromide (PTMD90-0015, 11): A mixture of 9 (108 µL, 229 mg, 1.00 mmol, 1.0 equiv) and 10 (352 µL, 325 mg, 2.40 mmol, 2.4 equiv) was stirred at 145 °C for 2 h. The reaction mixture was concentrated in vacuo and the residue was purified by recrystallization from EtOAc/EtOH (1:2) to afford 11 (196 mg, 40%) as a colorless solid. m.p. 248 °C; ^1^H NMR (400 MHz, CD_3_OD): δ = 1.45 (s, 18 H), 4.43-4.69 (m, 3 H), 4.97-5.22 (m, 2 H), 8.11-8.24 (m, 4 H), 8.83-9.01 (m, 4 H); ^13^C NMR (101 MHz, CD_3_OD): δ = 30.19, 37.70, 63.74, 71.06, 126.42, 146.09, 173.58; IR (film): L = 2972, 1643, 1468, 1117 cm^-1^; HRMS (ESI): *m/z* calcd for C_21_H_32_N_2_OBr_2_-H^+^-2Br^-^: 327.2436 [*M*-Br-Br-H]^+^; found: 327.2430.

### 2.2 UNC0642 MS Binding Assays

Competitive MS Binding Assays were performed as previously described (Kaiser et al., 2024; Nitsche et al., 2024) apart from one minor difference: in order to obtain competition curves for compounds bearing a 4-aminopyridinium moiety, the non-linear regression function “log(inhibitor) vs. normalized response – Variable slope” (Prism Software v. 6.07, GraphPad Software, La Jolla, CA, USA) was used instead of the recently used function “One site – fit Ki”. Statistically significant differences were verified by a two-sided *t*-test (alpha = 0.05).

### 2.3 Rat diaphragm myography

All procedures using animals followed animal care regulations. Preparation of rat diaphragm hemispheres from male Wistar rats (300 ± 50 g) and experimental protocol of myography was performed as previously described (Nitsche et al., 2024; Seeger et al., 2012). In brief, for all procedures (including wash-out steps, preparation of soman and test compound solutions) aerated Tyrode solution (125 mM NaCl, 24 mM NaHCO_3_, 5.4 mM KCl, 1 mM MgCl_2_, 1.8 mM CaCl_2_, 10 mM glucose, 95% O_2_, 5% CO_2_; pH 7.4; 25 ± 0.5 °C) was used. After the recording of control muscle force one hour after preparation, the muscles were incubated in the Tyrode solution, containing 3 μM soman for 20 min. Following a 20 min wash-out period, the test compounds were added in ascending concentrations (0.1 μM to 300 μM). The incubation time was 20 min for each concentration. The electric field stimulation was performed with 10 μs pulse width and 2 A amplitudes. The titanic stimulation of 20 Hz, 50 Hz, 100 Hz were applied for 1 s and in 10 min intervals. Measurements on non-toxic muscle were carried out according to the same scheme. Instead of soman, pure Tyrode was incubated. Muscle force was calculated as a time-force integral (area under the curve, AUC) and constrained to values obtained for maximum force generation at the start of the measurements (muscle force in the presence of Tyrode solution without any additives; 100%). All results were expressed in means ± SD (n = 6 - 12). Prism 5.0 (GraphPad Software, San Diego, CA, USA) was used for data analysis.

### 2.4 MD simulations

The model of the human muscle type nAChR was generated using Modeller with the PDB structure of the α7-nAChR as the template (PDB ID: 7KOX (Noviello et al., 2021). The orthosteric ligand nicotine and a sodium ion in the transmembrane pore were included by aligning the structure of the α3β4-nAChR (PDB ID: 6PV7 (Gharpure et al., 2019) to the homology model. Nicotine was subsequently minimized using SZYBKI (OpenEye Scientific Software, 2021). MB327 was docked in MB327-PAM-1 using MOE with an induced-fit refinement using default parameters (Chemical Computing Group, 2020). Ligand charges were calculated using Gaussian16 (M. J. Frisch et al., 2016) at the HF 6-31G* level of theory; force field parameters for the ligand were taken from the gaff force field (Wang et al., 2004). Using Packmol-Memgen (Schott-Verdugo and Gohlke, 2019), the system was embedded in a membrane containing 1-palmitoyl-2-oleoyl-*sn*-glycero-3-phosphocholine (POPC) lipids, solvated using the Optimal Point Charge water model (Izadi et al., 2014) with a minimum distance of 12 Å between receptor atoms and the edge of the box, KCl was added in a concentration of 150 mM, and the system was neutralized using Cl^-^ ions. To perform MD simulations, the AMBER22 package of molecular simulations software (Case et al., 2005; Case et al., 2022), the ff19SB force field (Tian et al., 2020) for the protein, the Lipid17 force field (Gould et al., unpublished) for lipids, the gaff force field for the ligand, and the Joung and Cheatham parameters for monovalent ions were used (Joung and Cheatham, 2008). MD simulations were performed as described previously (Kaiser et al., 2023). In short, a combination of steepest descent and conjugate gradient minimization was performed while gradually decreasing positional harmonic restraints. The system was then heated in a stepwise manner to 300 K, and harmonic restraints on receptor and ligand atoms were gradually removed subsequently. Then, 12 replicas of 1 μs long unbiased MD simulations were performed in the NPT ensemble using semi-isotropic pressure adaptation with the Berendsen barostat. The RMSD, electron density profiles and representative binding modes were computed using CPPTRAJ (Roe and Cheatham, 2013), as implemented in AmberTools (Case et al., 2023).

### 2.5 GIST computations

GIST (Lazaridis, 1998; Nguyen et al., 2011; Nguyen et al., 2012; Ramsey et al., 2016) computations, as implemented in CPPTRAJ (Roe and Cheatham, 2013), were performed in replicas where MB327 remained in the binding site (distance to I64δ (respectively, I61ε, I61α, I64β) < 5 Å, as done previously (Kaiser et al., 2023) during MD simulations. The backbone of each frame during these MD simulations was aligned to the starting structure of the simulations using CPPTRAJ (Roe and Cheatham, 2013). The middle carbon atom of the C3-linker in MB327 was used as the center of the box for GIST grid generations with grid dimensions of 40 increments along each axis and a grid spacing of 0.5 Å. The results were filtered based on the density of oxygen centers in each voxel compared to bulk density (cutoff: > 1.75) and the solvent free energy (cutoff: > −0.25 kcal mol^-1^). Visualization of water clusters and manual modification of MB327 to PTMD90-0012 (**8**) and PTMD90-0015 (**11**) was performed using MOE (Chemical Computing Group, 2020).

### 2.6 Image generation

Images of nAChR in complex with MB327 and its analogs were created using UCSF Chimera (Pettersen et al., 2004).

## 3 Results and Discussion

### 3.1 Synthesis of novel MB327 analogs

To identify novel non-symmetric MB327 analogs that should exhibit higher binding affinities to the MB327-PAM-1 binding site as well as higher intrinsic activities compared to MB327, a series of non-symmetric bispyridinium compounds **6a**-**6i** with a 4-aminopyridinium ion partial structure derived from compounds **1**-**3** was synthesized. In addition, based on modeling studies (see chapter 3.3), novel MB327 analogs **8** and **11** with an additional OH function were synthesized with the assumption that this modification should increase the binding affinity by displacing water molecules from the binding pocket.

#### Non-symmetric MB327 analogs PTM0064-PTM0072 (**6a**-**6i**) and PTMD90-0012 (**8**)

Non-symmetric MB327 analogs **6a**-**6h** bearing a 4-aminopyridinium ion moiety were readily accessible in one step by *N*-alkylation of 4-aminopyridines **5** with *N*-(3-iodopropyl)pyridinium building block **4**, analogous to the method described by Rappenglück *et al*. (Rappenglück et al., 2018) (Scheme 1). To cover a wider variety of 4-amino substituents in the target compounds, the set of commercially available 4-aminopyridines **5a**, **5b** and **5e**-**5g** was extended with some building blocks (**5c**, **5d** and **5h**), synthesized according to literature (Hay et al., 2015; Price et al., 2006; Wang et al., 2019). *N*-Alkylation of the pyridines **5a** and **5c**-**5h** with building block **4** (Rappenglück et al., 2018) was accomplished by stirring the components in acetonitrile at 90 °C under microwave irradiation for 1 h. After removing the reaction solvent, the resulting residues were purified by crystallization to yield the target compounds **6a** and **6c**-**6h** in good to excellent yields (72-93%) and high purities (≥ 98%). Reaction with *tert*-butyl pyridin-4-ylcarbamate (**5b**), however, required a lower reaction temperature to prevent cleavage of the Boc group as a reaction at 90 °C for 1 h had led to a mixture of product **6b** and 4-amino-analog **1** in a ratio of 3:1. By stirring **5b** with building block **4** at 60 °C for 15 h no side product **1** was observed and the product **6b** was afforded in excellent yield (92%) and sufficient purity (94%). To get the hydroiodide **6i**, the Boc protecting group of **6h** was cleaved by stirring with trimethylsilyl iodide (TMSI) (4.0 equiv) in acetonitrile at room temperature for 1 h (Lott et al., 1979). Thus, compound **6i** was isolated in quantitative yield (99%) and with high purity (100%).

**Scheme 1.**
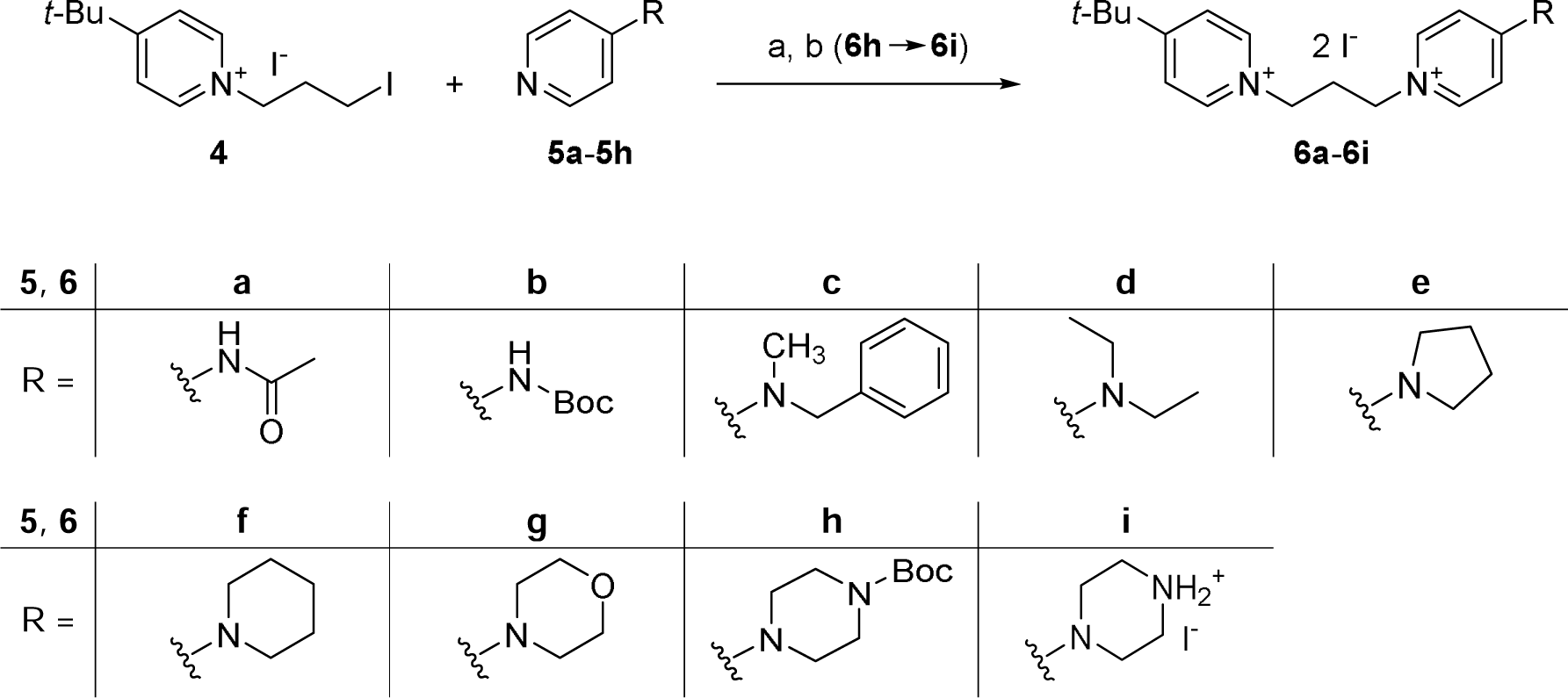
Synthesis of **6a-6i**. Reagents and conditions: (a) 4-(*tert*-butyl)-1-(3-iodopropyl)pyridin-1-ium iodide (**4**) (1.0 equiv), pyridines **5a**-**5h** (1.05-1.1 equiv), MeCN, microwave: 150 W, 60-90 °C, 1-15 h, 72-93%; (b) **6i**: **6h** (1.0 equiv), TMSI (4.0 equiv), MeCN, rt, 1 h, 99%.

PTMD90-0012 (**8**) was synthesized from building block **4** (Rappenglück et al., 2018) and 7-hydroxyquinoline (**7**) under the reaction conditions described for bispyridinium compounds **6a**-**6h**. However, the reaction time had to be increased to 3 h to compensate for the lower reactivity of the sterically more demanding quinoline **7** as compared to the 4-aminopyridines **5**. That way, PTMD90-0012 (**8**) was obtained in 29% yield after recrystallization (purity ≥ 98%) (Scheme 2).

**Scheme 2.**
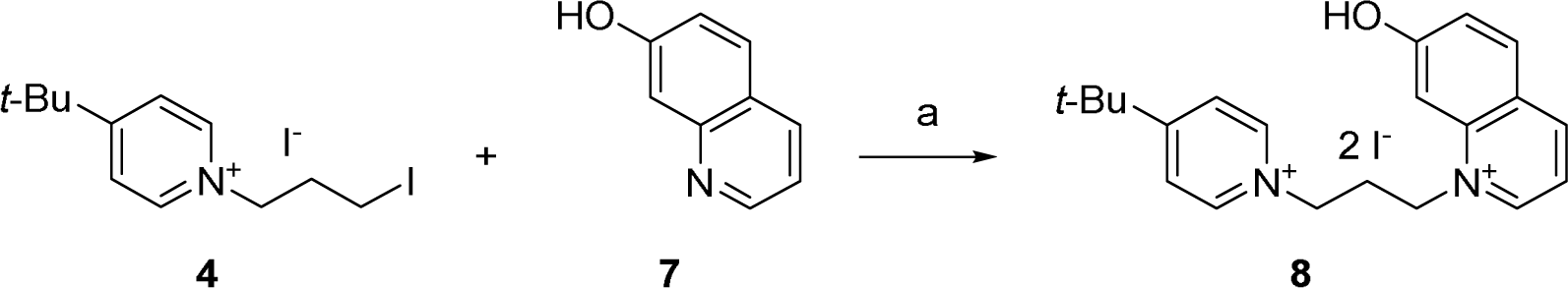
Synthesis of PTMD90-0012 (**8**). Reagents and conditions: (a) 4-(*tert*-butyl)-1-(3-iodopropyl)pyridin-1-ium iodide (**4**) (1.0 equiv), 7-hydroxyquinoline (**7**) (1.1 equiv), MeCN, microwave: 150 W, 90 °C, 3 h, 29%.

#### MB327 analog PTMD90-0015 (**11**)

The symmetric bispyridinium compound PTMD90-0015 (**11**) with a 2-hydroxypropyl linker between the two pyridinium rings, was synthesized by heating 1,3-dibromopropan-2-ol (**9**) with an excess of 4-*tert*-butylpyridine (**10**) to 145 °C for 2 h. Bis-alkylation product PTMD90-0015 (**11**) was obtained in 40% yield and in high purity (100%) after recrystallization (Scheme 3). In contrast, reaction of **8** with epichlorohydrine or 1,3-dichloropropan-2-ol under the reaction conditions described above led only to the corresponding monosubstituted products.

**Scheme 3.**
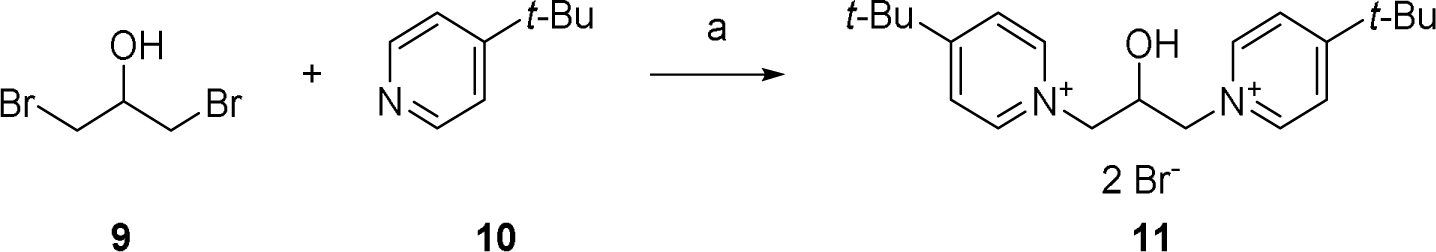
Synthesis of PTMD90-0015 (**11**). Reagents and conditions: (a) 1,3-dibromopropan-2-ol (**9**) (1.0 equiv), 4-*tert*-butylpyridine (**10**) (2.4 equiv), 145 °C, 2 h, 40%.

### 3.2 Biological evaluation

#### Affinity to the MB327-PAM-1 binding site of Torpedo californica nAChR

All of the newly developed compounds presented in this study, i.e. PTM0064-PTM0072 (**6a**-**6i**), PTMD90-0012 (**8**) and PTMD90-0015 (**11**) as well as MB327 and its recently reported analogs PTM0062 (**1**) (Kaiser et al., 2023), PTM0063 (**2**) (Kaiser et al., 2023) and PTM0056 (**3**) (Rappenglück et al., 2018) were evaluated for their binding affinity towards the MB327-PAM-1 binding site of *Torpedo californica* nAChR by means of the recently introduced UNC0642 MS Binding Assays (Table 1) (Kaiser et al., 2024; Nitsche et al., 2024). First, for economic reasons, all test compounds were studied at a single concentration of 10 µM and a reporter ligand concentration of 1 µM. For a set of selected compounds also the binding affinity constants (p*K*_i_ values) were determined in full-scale MS competition experiments.

**Table 1.**
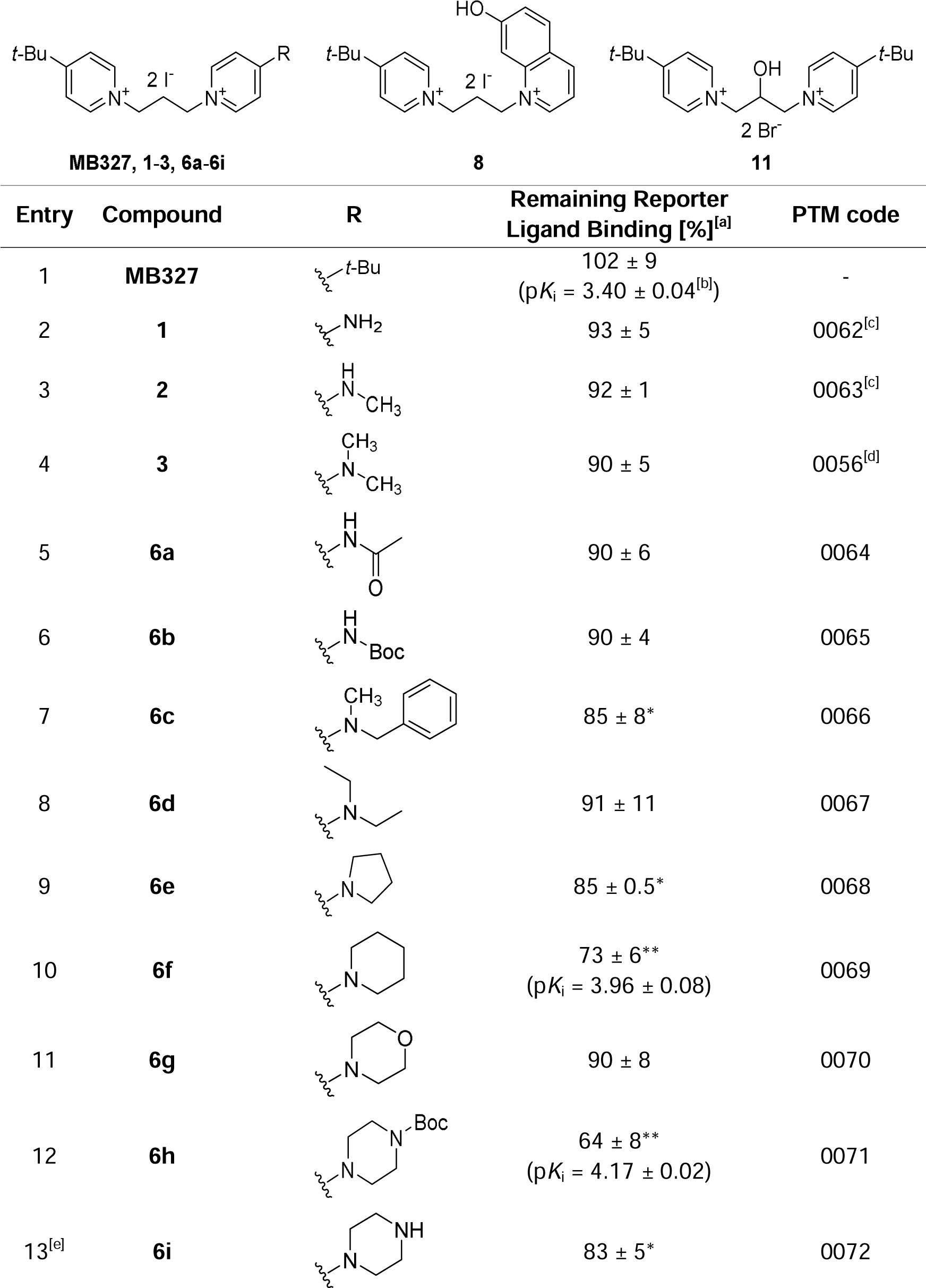
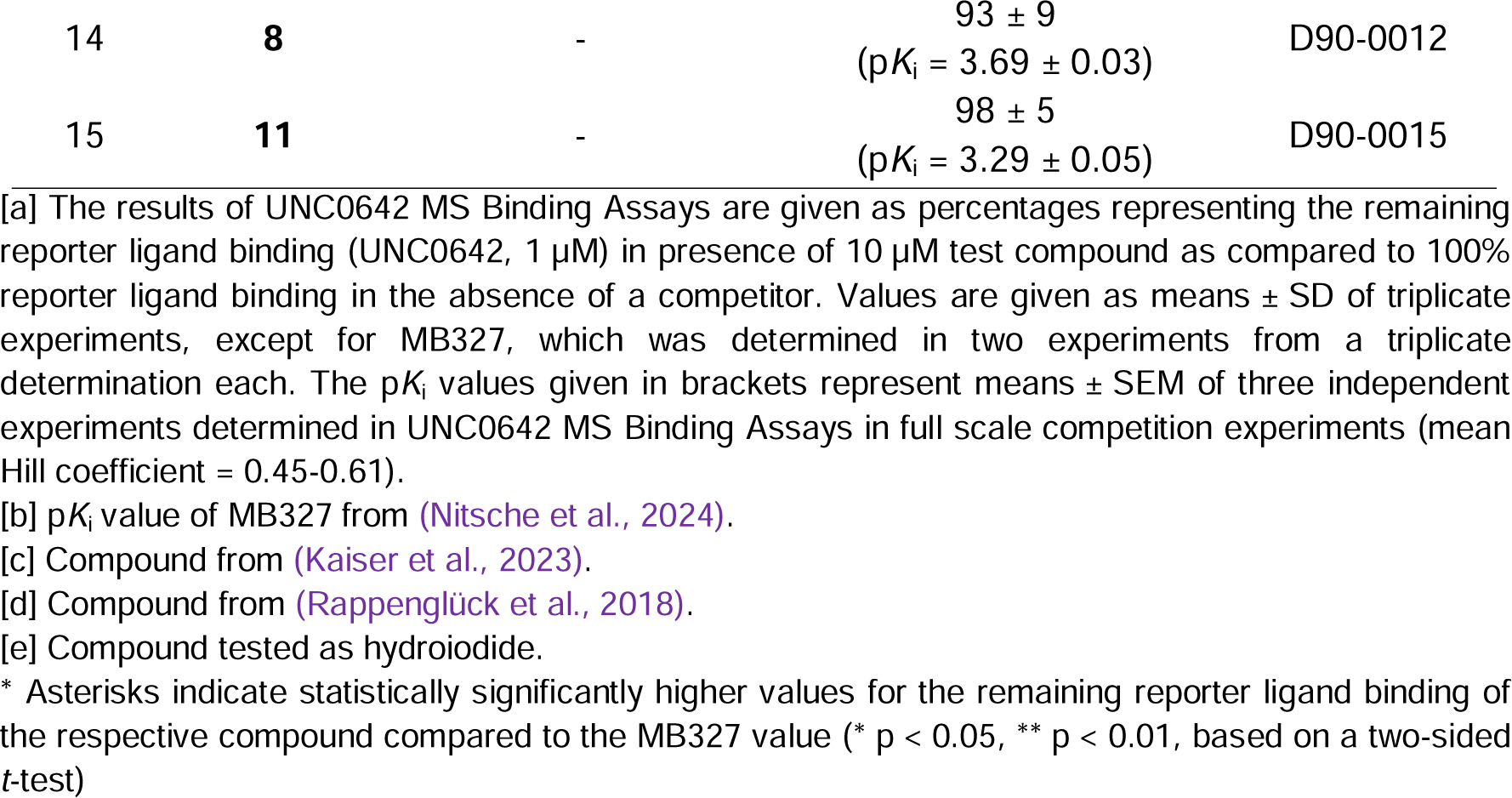
Binding affinities of bispyridinium compounds for the MB327-PAM-1 binding site of *Torpedo californica* nAChR, determined in UNC0642 MS Binding Assays.

Residual reporter ligand binding in the presence of 10 µM of the respective compounds listed in Table 1 range from 102% ± 9% for MB327 (Table 1, entry 1) to 64% ± 8% for the *N-*Boc-piperazino derivative PTM0071 (**6h**, Table 1, entry 12). Substitution of one of the two 4-*tert*-butyl residues of MB327 by an amino [PTM0062 (**1**), 93% ± 5%, Table 1, entry 2], an *N*-methylamino [PTM0063 (**2**), 92% ± 1%, Table 1, entry 3], or dimethylamino group [PTM0056 (**3**), 90% ± 5%, Table 1, entry 4] results in a nominal yet not statistically significant reduction of residual reporter ligand binding as compared to MB327. This is also true for compounds PTM0064 (**6a**, 90% ± 6%, Table 1, entry 5) and PTM0065 (**6b**, 90% ± 4%, Table 1, entry 6) displaying an *N*-acetamido- and an *N*-*tert*-butoxycarbonylamino moiety, respectively effecting a reduction to 90% of remaining reporter ligand binding. According to the data obtained for **1**-**3** and **6a**-**6b** the capability of the nitrogen substituents, present in these compounds to participate in hydrogen bonding seems to have only little if any influence on the binding affinities, the remaining reporter ligand binding amounting in any case to about 90%. Likewise, when the dimethylamino group in PTM0056 (**3**, 90% ± 5%, Table 1, entry 4) is replaced by a sterically more demanding *N*,*N*-diethylamino moiety, the binding affinity of the resulting PTM0067 (**6d**) remains with 91% ± 11% virtually unaltered (remaining reporter ligand binding, Table 1, entry 8). Although **1**-**3** as well as **6a**, **6b** and **6d** effect remaining reporter ligand binding nominally below that of MB327 (Table 1, entry 1), for none of these compounds the observed differences are statistically significant. However, for the *N*-benzyl-*N*-methylamino derivative PTM0066 (**6c**) with an enlarged lipophilic domain, remaining reporter ligand binding reaches 85% ± 8% (Table 1, entry 7), which is statistically significantly lower than that of MB327 (102% ± 9%, Table 1, entry 1).

Notably, the pyrrolidino and the piperidino substituted derivatives PTM0068 (**6e**) and PTM0069 (**6f**), of which the former can be considered as a cyclic analog of the diethylamino substituted **6d,** reduce remaining reporter ligand binding to 85% ± 0.5% (Table 1, entry 9) and 73% ± 6% (Table 1, entry 10), both values being also statistically significantly below than that of MB327 (102% ± 9%, Table 1, entry 1).

Upon transition from the piperidino-substituted bispyridinium salt **6f** to the more polar morpholino-substituted analog PTM0070 (**6g**), again a decline in binding affinity is observed with the remaining reporter ligand binding increasing to 90% ± 8% (Table 1, entry 11). However, for the nitrogen analog of the morpholino derivative **6g**, the piperazino derivative PTM0072 (**6i**), the decline in binding affinity compared to **6f** displaying a piperidino residue (73% ± 6%, Table 1, entry 10) was less pronounced, with the remaining reporter ligand binding amounting to 83% ± 5% (Table 1, entry 13). Obviously, the binding affinity-diminishing effect of the additional heteroatom seems to be less distinct for the piperazino derivative **6i** than for the morpholino derivative **6g** (as compared to the piperidino analog **6f**), which can be possibly attributed to the capability **6i** to act as a hydrogen bridge acceptor and donor.

Remarkably, the hydrogen bridge donor capability appears to be of minor importance, as upon attachment of an *N*-Boc substituent to the piperazino moiety of **6i** resulting in PTM0071 (**6h**, Table 1, entry 12), the binding affinity is distinctly improved despite the absence of the formerly present NH group. Thus, a remaining reporter ligand binding of 64% ± 8% could be measured for this compound, which is statistically significantly lower than that of the piperazino-substituted compound **6i** (Table 1, entry 13). Hence, the piperidino-substituted compound **6f** (Table 1, entry 10) and the *N*-Boc-piperazino derivative **6h** (Table 1, entry 12) possess the highest binding affinities for the MB327-PAM-1 binding site of the nAChR of the compounds evaluated in this study.

Modeling studies (see chapter 3.3) indicated, that the presence of an OH function in ligands of the MB327-PAM-1 binding site might be favorable for the binding affinity by displacing water molecules present in the binding pocket. In particular, the MB327 analog **11** with an OH function in the spacer linking the two pyridinium subunits in the molecule as well as **8** with a 7-hydroxyquinolinium replacing one of the two pyridinium subunits in MB327 were expected to possess improved binding affinities. Hence, the binding affinities of PTMD90-0015 (**11**) and PTMD90-0012 (**8**) were studied. Both compounds, however, show no or only a negligible improvement of the binding affinity compared to MB327 (Table 1, entry 1), the remaining reporter ligand binding amounting to 98% ± 5% for **11** (Table 1, entry 15) and 93% ± 9% for **8** (Table 1, entry 14). Although no clear improvement of the binding affinity could be achieved with **11** and **8**, the results indicate that the presence of an additional polar OH function is tolerated in the binding pocket.

Finally, for a small set of compounds, the p*K*_i_ values as a more accurate measure of the binding affinity were determined in full-scale competitive MS Binding Assays. This set comprised the two compounds **6f** and **6h**, which had shown the highest binding affinities in the single point determinations described above (reduction of the remaining reporter ligand binding to < 75%) as well as the two OH function-containing derivatives **8** and **11,** which had emerged as candidate compounds for testing from modeling studies. Notably, the piperidino derivative **6f** (Table 1, entry 10) exhibits a p*K*_i_ of 3.96 ± 0.08, which was approximately 0.5 log units and thus statistically significantly higher than the p*K*_i_ of MB327 previously determined to be 3.40 ± 0.04 (Table 1, entry 1) (Nitsche et al., 2024). The *N*-Boc-piperazino derivative **6h** (Table 1, entry 12) showed an even higher p*K*_i_ of 4.17 ± 0.02, as to be expected from the preliminary binding data (remaining reporter ligand binding 64% ± 5%), exceeding that of MB327 by approximately 0.8 log units. For the MB327 analog **11** with an OH function as part of the propan-1,3-diyl spacer (Table 1, entry 15), the p*K*_i_ value (3.29 ± 0.05) matched that of MB327 within the limits of error. For the 7-hydroxyquinolinium derivative **8** (Table 1, entry 14), a p*K*_i_ value of 3.69 ± 0.03 was found, almost 0.3 log units higher than that of MB327, which is statistically significantly different, indicating the higher binding affinity of this compound compared to MB327.

#### Evaluation of muscle force recovery in soman-poisoned rat diaphragm hemispheres

In addition to the binding affinities, the intrinsic activities of a selection of the test compounds have been determined by means of an *ex vivo* assay based on rat diaphragm hemispheres. In this assay dissected rat diaphragm hemispheres are treated with a solution containing 3 µM soman. Whereas upon indirect electric field stimulation commonly carried out at frequencies of 20 Hz, 50 Hz and 100 Hz, unpoisoned rat diaphragms undergo muscle contractions, in case of poisoned samples no or only very faint contractions occur. This inhibition further does not vanish, when the poisoned samples are freed from the toxin by washing, typically performed as a control, as this has no effect on the irreversible inhibition of the AChE by the nerve agent. The positive intrinsic activity of the test compounds becomes apparent, when their addition to the poisoned muscle preparations, typically performed in increasing concentrations from 0.1-300 µM, affects a recovery of the muscle force of the soman-impaired rat diaphragm hemispheres. Muscle force inhibition reappears, when samples are subjected to a subsequent washout step, which is due to the irreversible inactivation of the AChE and indicative of the reversibility of the receptor-mediated resensitizing effect of the test compounds.

For the characterization of the muscle force restoring potency of soman-poisoned rat diaphragms in the aforementioned test system, the piperidino- and the *N*-Boc-piperazino derivatives **6f** and **6h** were selected as they had shown the highest binding affinities among the new test compounds presented in this study. In addition, the piperazino derivative **6i**, closely related to **6h**, the *N*,*N*-dimethylamino-substituted compound **3** as parent compound, as well as the hydroxy-substituted compounds **8** and **11** as analogs of MB327 devoid of an amino-substituent were evaluated in this test system.

Since it is known, that the largest efficacy is observed at low stimulation frequencies (Seeger et al., 2012), only the results of experiments performed at 20 Hz will be presented (Figure 2) and discussed in the following. The data of measurements carried out at higher frequencies (50 Hz, 100 Hz) can be found in the Supporting Information (SI, Figure S2).

**Figure 2:**
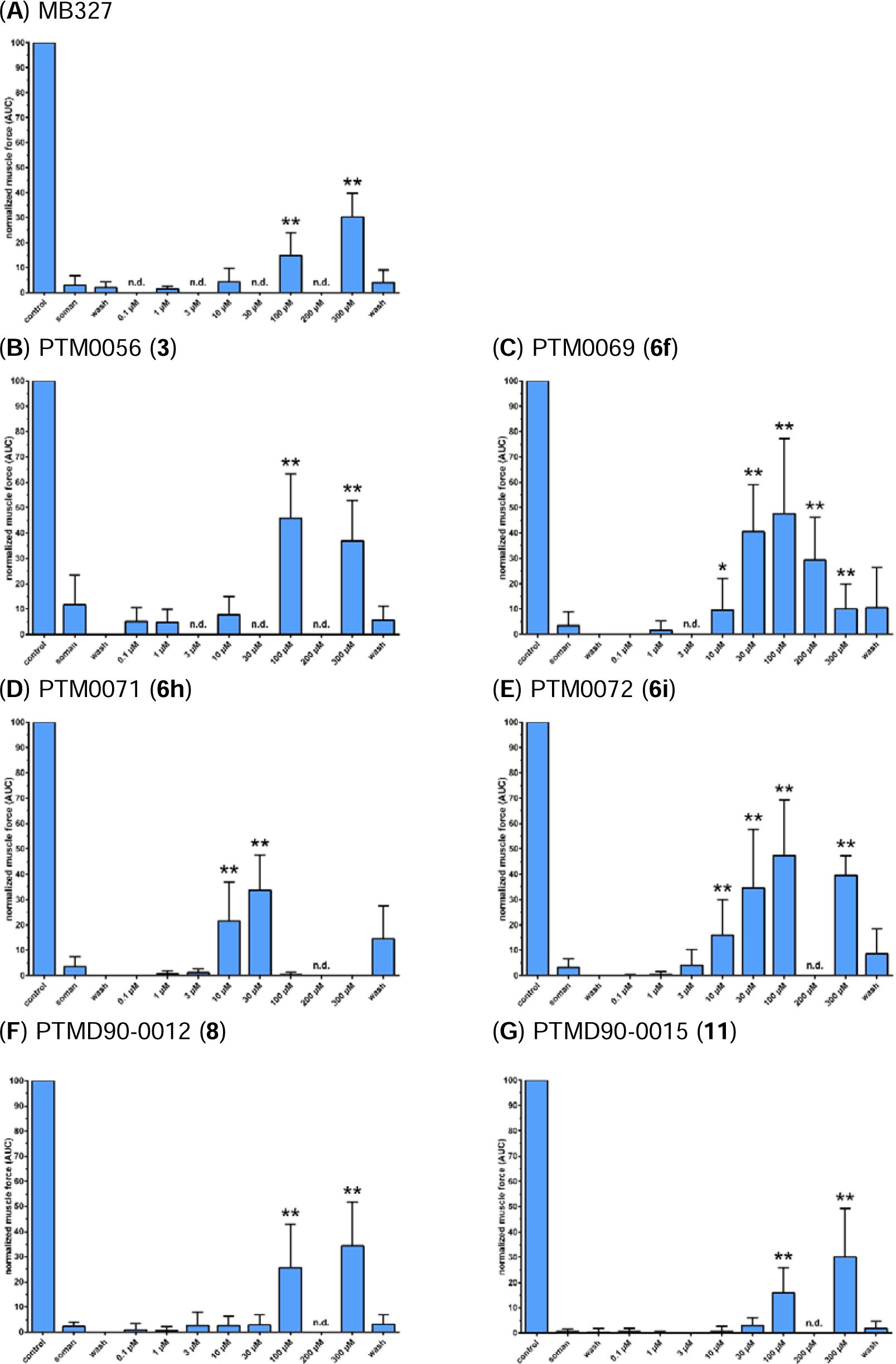
Concentration-dependent restoration of muscle force of soman-poisoned rat diaphragms by (**A**) MB327 (Niessen et al., 2018), (**B**) PTM0056 (**3**), (**C**) PTM0069 (**6f**), (**D**) PTM0071 (**6h**), (**E**) PTM0072 (**6i**), (**F**) PTMD90-0012 (**8**) and (**G**) PTMD90-0015 (**11**). For indirect stimulation, a frequency of 20 Hz was applied. Data are shown as % of control and are given as mean ± SD (n = 4-25). Asterisks indicate statistically significantly higher values for the respective compound concentration compared to the soman value (* p < 0.05, ** p < 0.01, based on a two-sided *t*-test). n.d.: not determined.

As reference, the published data obtained for MB327 in the test system have been included in Figure 2 (Niessen et al., 2018). As previously reported, MB327 addition (Figure 2A) leads to a concentration-dependent reactivation of soman-impaired muscle. The recovery gradually increases with the MB327 concentration (Figure 2A). At a MB327 concentration of 100 µM, the muscle force recovery amounts to 14.8% ± 9.2%, reaches 30.2% ± 9.5% at 300 µM (both values statistically significantly different from the value obtained for the soman-poisoned muscle), to finally decrease to 4.2% ± 4.2% at 1000 µM MB327 (not shown in Figure 2A)(Niessen et al., 2018).

Similar to MB327, the analogs **3**, **6f**, **6h** and **6i** exhibiting a 4-amino group at one of the two pyridinium subunits instead of a *tert*-butyl moiety (present in MB327) lead to an increase of the regeneration of the muscle force of soman-poisoned rat diaphragms with increasing concentrations, whereas the onset and the size of this effect differ. The *N*,*N*-dimethylamino-substituted analog **3** shows its maximum recovery at a concentration of 100 µM (45.8% ± 17.4%), which statistically significantly exceeds that of MB327 (p < 0.01) at the same concentration (14.8% ± 9.2%). However, the reactivation decreases nominally at 300 µM to 37.0% ± 15.8%, approximating that of MB327 at the same concentration (30.2% ± 9.5%). At a compound concentration of 100 µM, the results for the piperidino- and the piperazino-substituted compounds **6f** (47.5% ± 29.8%) and **6i** (47.3% ± 21.8%) are comparable to the value observed for the dimethylamino derivative **3**, which are also nominally the highest values of all three compounds. At concentrations above 100 µM, alike observed for the dimethylamino derivative **3**, the muscle force declines again for **6f** as well as **6i**, which is in the case of **6i** with 39.5% ± 7.7% muscle force recovery at 300 µM compound concentration far less pronounced than for **6f** with 10.0% ± 9.8% (statistically significant, p < 0.01%; **6f** at 200 µM 29.3% ± 16.8%). The *N*-Boc-piperazino derivative **6h** induces a rather strong muscle force recovery already at 10 µM with a value amounting to 21.7% ± 15.3%, which is nominally the highest for all tested compounds at this concentration, which further increases at 30 µM to 33.8% ± 13.8% followed by a sharp drop to almost 0% (0.3% ± 0.8%) at 100 µM (0.0% ± 0.0% at 300 µM). This strong muscle force recovery effect of **6h** at a low concentration might be a result of its high binding affinity, which is the highest of all compounds studied.

Notably, all three MB327 analogs with cyclic amino residues exert already at a concentration of 10 µM a muscle force recovery that is statistically significantly higher than that of the soman-poisoned but untreated muscle, whereas this is neither the case for the dimethylamino derivative **3** nor for MB327. For the piperidino derivative **6f,** the respective recovery of muscle force at 10 μM amounts to 9.5% ± 12.7%, for the *N*-Boc-piperazino analog **6h** to 21.7% ± 15.3% and the piperazino derivative **6i** to 16.0% ± 14.1%. Statistical significance of muscle force recovery over the soman-poisoned but untreated muscle as reference value is reached for these three compounds also at a test compound concentration of 30 µM (**6f**, 40.6% ± 18.5%; **6h**, 33.8% ± 13.8%; **6i**, 34.5% ± 23.2%). Yet, it remains unclear, whether this is also the case for the dimethylamino derivative **3** and MB327, as data for these two compounds when applied at 30 µM are missing.

The two MB327 analogs featuring an additional hydroxy group either in the spacer, **11**, or as part of a quinolinium moiety, **8**, show very similar effects in terms of muscle force regeneration after soman-intoxication. Also, muscle force recovery effected by **11** and **8** at concentrations of 100 μM and 300 μM is statistically significant as compared to the soman value. Thereby, the values for muscle force recovery of soman-poisoned muscle amounted for **11** to 15.9% ± 9.9% at 100 μM and to 30.1% ± 19.1% at 300 μM, whereas those for **8** are higher with 25.6% ± 17.5% at 100 µM and 34.3% ± 17.5% at 300 µM. For both compounds, no decrease in muscle force could be observed up to the highest concentration of 300 µM studied.

Finally, the effects of test compounds that were still available in sufficient amounts, i.e., of MB327 and the dimethylamino **3**, the piperidino **6f** as well as the quinolinium derivative **8,** on the muscle force of rat diaphragm hemispheres not been poisoned with soman were studied (SI, Figure S3). These experiments were carried out in analogy to those for the determination of muscle force recovery of soman-poisoned rat diaphragm hemispheres. Accordingly, the muscle force of rat diaphragm hemispheres not exposed to soman was measured as a function of increasing concentrations of the test compounds under indirect electric field stimulation at 20 Hz, 50 Hz and 100 Hz. Data from analogous experiments performed in parallel but without the application of test compounds served as reference by unveiling the loss of muscle force purely due to the washing steps. MB327 and dimethylamino derivative **3** showed no inhibitory effect up to the maximum concentration of 300 µM, as did the quinolinium derivative **8** up to 200 µM as the highest concentration still feasible due to a shortage of the compound. For the piperidino derivative **6f**, however, although the muscle force remained unaffected up to a concentration of 100 µM, a distinct reduction occurred at a concentration of 300 µM. The latter might explain the bell-shaped curve observed for this compound, **6f**, in the muscle force recovery experiments of soman-poisoned rat diaphragm hemispheres (Figure 2C). The positive effect of **6f** on soman-poisoned rat diaphragms mediated by binding to the MB327-PAM-1 binding site of the nAChR might be counteracted by a direct negative effect on muscle force, mediated by different binding sites, which seems to become dominating at concentrations above 100 µM of **6f,** leading to the observed decline of muscle force recovery in soman-poisoned muscles (Figure 2C) above this concentration.

### 3.3 Substituting energetically unfavorable water clusters using *in silico* methods

We performed unbiased MD simulations (12 replicas of 1 µs length each) of the human nAChR with MB327 bound to the recently identified allosteric binding site MB327-PAM-1 in all five subunits (Kaiser et al., 2023). During the simulations, the receptor and membrane stayed structurally invariant shortly after the simulations started (SI, Figure S4, S5). MB327 mostly or completely remained within the binding sites in between the α- and δ-subunits (ten out of twelve replicas) as well as in between the α- and ε-subunits (twelve out of twelve replicas) using the distance of heavy atoms of MB327 to heavy atoms of I64δ (respectively I61ε) to characterize unbinding as done previously (Kaiser et al., 2023). For these two binding sites, Grid Inhomogeneous Solvent Theory (GIST) computations (Lazaridis, 1998; Nguyen et al., 2011; Nguyen et al., 2012; Ramsey et al., 2016) were subsequently performed to identify potential energetically unfavorable water clusters (Figure 3B, SI, Figure S6A). We then visualized the docked MB327 binding mode and representative MB327 structures from MD simulations in the presence of such water clusters and manually created new MB327 analogs with substituents replacing these water clusters. Initially, during the first few replicas (5 out of 12) of MD simulations and subsequent GIST computations, a network of energetically unfavorable water molecules near the linker of MB327 was identified between E65α, V66α, and Q68α (SI, Figure S7). Based on these preliminary results, PTMD90-0015 (**11**) was designed. However, after completion of the MD simulations, the hydroxyl group located at the C3-linker would not result in replacing energetically unfavorable water molecules as indicated by GIST. Hence, PTMD90-0012 (**8**) was designed based on the results after 1 µs long simulations (Figure 3C, SI, Figure S6B), such that it should substitute energetically unfavorable water molecules in close proximity to I63ε, E65ε, and E204ε (respectively L66δ, E68δ, and E210δ). As only the analogs based on the docked structure [PTMD90-0012 (**8**)] and those based on the initial MD simulations results [PTMD90-0015 (**11**)] could be synthesized, they were subsequently tested for affinity and resensitizing potential.

**Figure 3:**
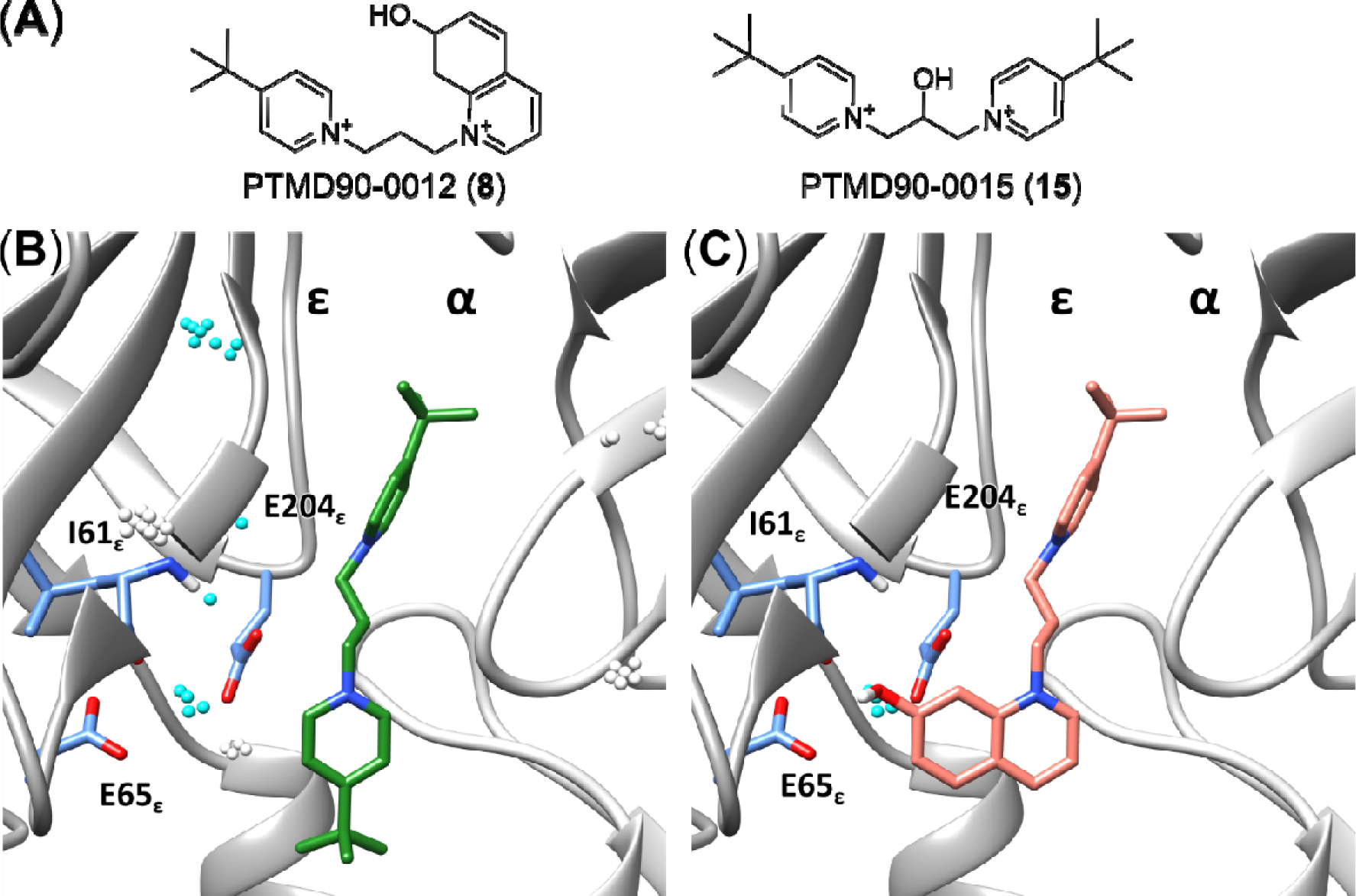
Design of MB327 analogs based on substituting water clusters in MB327-PAM-1. **A)** Structure of the MB327 analogs PTMD90-0012 (**8**) and PTMD90-0015 (**11**). **B)** MB327 (green) binding in between the α- and ε-subunit. Water clusters identified during GIST computations are shown as spheres. Water clusters within 5 Å of MB327 are shown in cyan. **C)** Proposed binding mode of PTMD90-0012 (**8**) (salmon) in between the α- and ε-subunit. Modification of MB327 to PTMD90-0012 (**8**) (salmon) leads to a substitution of a water cluster located in proximity to I63_ε_, E65_ε_, and E204_ε_; for I61_ε_, the backbone atoms are shown in addition to the side chain.

## 4 Conclusion

The recently developed non-symmetric MB327 analogs PTM0062 (**1**), PTM0063 (**2**) and PTM0056 (**3**), which have an amino, methylamino or dimethylamino group in place of one of the two *tert*-butyl residues of MB327, respectively, show higher muscle force restoring activity on soman-poisoned rat diaphragms than MB327. They are therefore promising starting points for the development of new resensitizers for desensitized muscle-type nAChRs as antidots for OPC poisoning.

In the present study, a series of new non-symmetric MB327 analogs, PTM0064-PTM0072 (**6a**-**6i**), with a 4-amino-substituted pyridinium ion substructure derived from compounds **1**-**3**, were synthesized and evaluated for their binding affinity towards the MB327-PAM-1 binding site of *Torpedo californica* nAChR using the recently introduced UNC0642 MS Binding Assays. In addition, selected compounds were evaluated for their intrinsic activity on soman-poisoned rat diaphragms in myography assays.

Among the compounds evaluated in this study, the piperidino derivative PTM0069 (**6f**) and the *N*-Boc-piperazino derivative PTM0071 (**6h**) showed the highest affinities for the MB327-PAM-1 binding site of the nAChR. The p*K*_i_ values for **6f** and **6h** exceeded that of MB327 by approximately 0.5 and 0.8 log units, respectively. PTMD90-0015 (**11**), designed to substitute unfavorable water clusters in MB327 PAM-1 after preliminary MD simulations, showed reduced affinity towards MB327-PAM-1 and no beneficial resensitizing effects compared to MB327. These results are concordant with that the hydroxy group located at the C3-linker would not result in replacing energetically unfavored water molecules as indicated by GIST after 1 µs long MD simulations. By contrast, PTMD90-0012 (**8**), designed using extended MD data, showed a statistically significant increase of affinity compared to MB327. Likely, this may result from replacing energetically unfavored water molecules close to I63_ε_, E65_ε_, and E204_ε_ (or L66_δ_, E68_δ_, and E210_δ_) with the 7-hydroxy quinazoline moiety.

*Ex vivo* studies in soman-poisoned rat diaphragms showed a clear muscle force restoring activity for all compounds tested [PTM0069 (**6f**), PTM0071 (**6h**), PTM0056 (**3**), PTM0072 (**6i**), PTMD90-0012 (**8**), and PTMD90-0015 (**11**)]. In particular, compounds PTM0069 (**6f**), PTM0071 (**6h**) and PTM0072 (**6i**), which all have a cyclic amino residue instead of one of the two *tert*-butyl residues of MB327, showed a statistically significantly higher activity at lower concentrations than MB327. Thus, to achieve muscle force recovery comparable to that of the aforementioned compounds, MB327 must be used at a tenfold higher concentration. This could be due to the fact that PTM0069 (**6f**) and PTM0071 (**6h**), in particular, but also PTM0072 (**6i**) have statistically significantly higher binding affinities for the MB327-PAM-1 binding site than MB327. It is noteworthy that the recovery of muscle force with PTM0069 (**6f**) and PTM0071 (**6h**) at higher concentrations decreased significantly immediately after reaching the maximum value, a phenomenon also observed for MB327. Experiments with PTM0069 (**6f**) and rat diaphragms that had not been poisoned with soman showed that in the presence of higher concentrations of PTM0069 (**6f**), a significant inhibition of muscle force occurs, which is reversible as it almost completely disappears after a subsequent washout step. Interestingly, this reversible inhibition occurs in the same concentration range in which the muscle force restoring effect of compound **6f** decreases in the experiments with soman-poisoned rat diaphragms.

Taken together, this indicates that the positive resensitizing effect mediated by binding to the MB327-PAM-1 binding site is counteracted by a muscle force inhibitory effect becoming generally prominent at higher concentrations that appears to be due to reversible binding to a different binding site.

Therefore, future efforts must focus, on the one hand, on further increasing the affinity of new compounds for the MB327-PAM-1 binding site and, on the other hand, on reducing the direct muscle inhibitory effect.

## Supporting Information

Supplementary data associated with this article can be found in the online version at doi: xxx

## 6. Acknowledgments

This work was supported by the German Ministry of Defence (E/ U2AD/KA019/IF558). We are grateful for computational support and infrastructure provided by the “Zentrum für Informations- und Medientechnologie” (ZIM) at the Heinrich Heine University Düsseldorf and the computing time provided by the John von Neumann Institute for Computing (NIC) to HG on the supercomputer JUWELS at Jülich Supercomputing Center (JSC) (user IDs: HKF7, VSK33, nAChR). HG is grateful to OpenEye Scientific Software for granting a Free Public Domain Research License.

