## Supplemental file for "Synthesis and Biological Evaluation of Novel MB327 Analogs as Resensitizers for Desensitized Nicotinic Acetylcholine Receptors after Intoxication with Nerve Agents"

[d]	Prof. Dr. H. Gohlke

John von Neumann Institute for Computing (NIC), Jülich Supercomputing Centre (JSC), Institute of Biological Information Processing (IBI-7: Structural Biochemistry) & Institute of Bio- and Geosciences (IBG-4: Bioinformatics)

Forschungszentrum Jülich

Wilhelm-Johnen-Strasse, 52428 Jülich (Germany)

**Table of Content**

Supplemental Figure S1 …………..……………….…………………………………………………………………….. 3

Supplemental Figure S2 …………..……………….…………………………………………………………………….. 4

Supplemental Figure S3 …………..……………….…………………………………………………………………….. 5

Supplemental Figure S4 …………..……………….…………………………………………………………………….. 6

Supplemental Figure S5 …………..……………….…………………………………………………………………….. 6

Supplemental Figure S6 …………..……………….…………………………………………………………………….. 7

Supplemental Figure S7 …………..……………….…………………………………………………………………….. 7

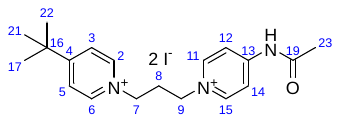

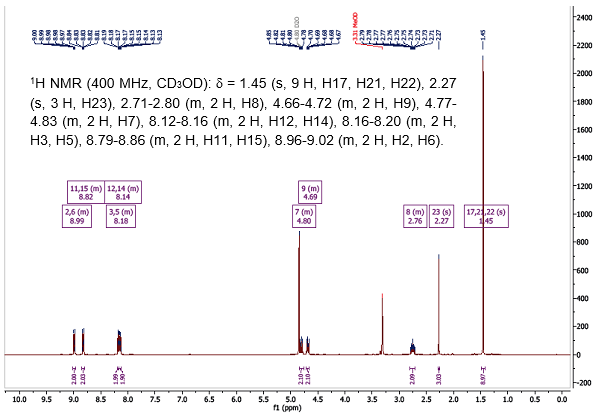

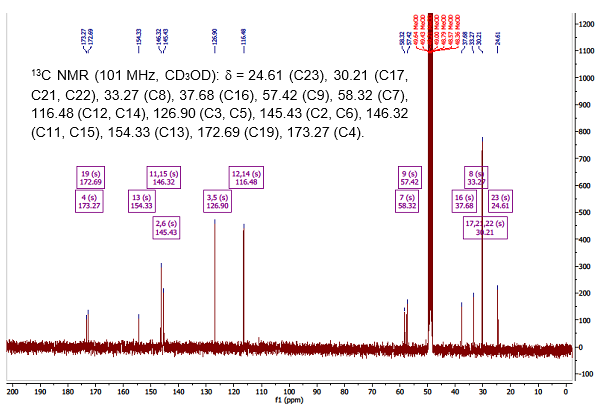

**Figure S1:** ^1^H and ^13^C NMR spectrum of PTM0064 (**6a**), as an example of a non-symmetric bispyridinium compound, with signals assigned.

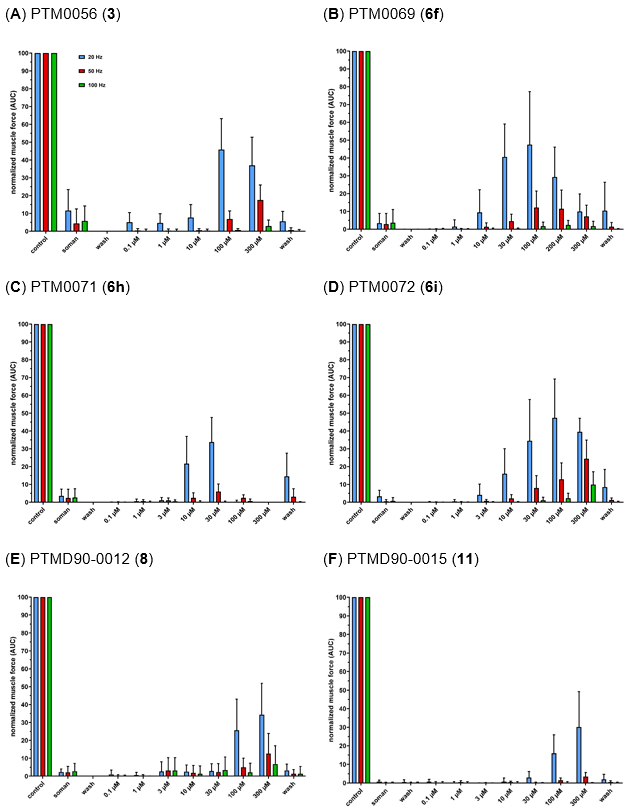

**Figure S2:** Concentration-dependent restoration of the muscle force by (**A**) PTM0056 (**3**), (**B**) PTM0069 (**6f**), (**C**) PTM0071 (**6h**), (**D**) PTM0072 (**6i**), (**E**) PTMD90-0012 (**8**), (**F**) PTMD90-0015 (**11**) of rat diaphragm preparations after poisoning with soman (3 µM). For indirect stimulation, a frequency of 20 Hz, 50 Hz and 100 Hz was applied. Data are shown as % of control and are given as mean ± SD (n = 4‑25).

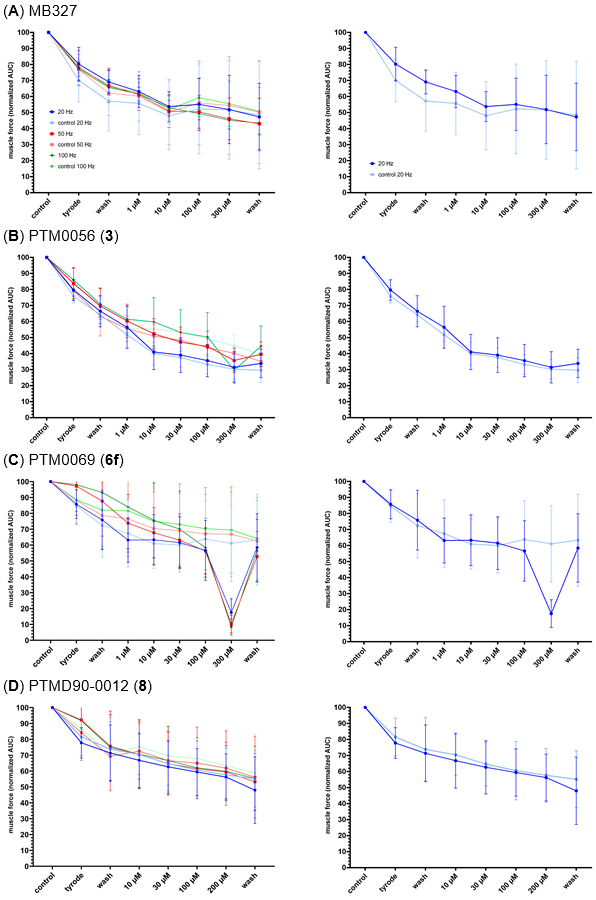

**Figure S3:** Muscle force of unpoisoned rat diaphragms after treatment with (**A**) MB327, (**B**) PTM0056 (**3**), (**C**) PTM0069 (**6f**) and (**D**) PTMD90-0012 (**8**). For indirect stimulation, a frequency of 20 Hz, 50 Hz and 100 Hz was applied. Muscle force generation was presented as the area under the curve normalized to muscle force under control conditions at the start of the measurement.

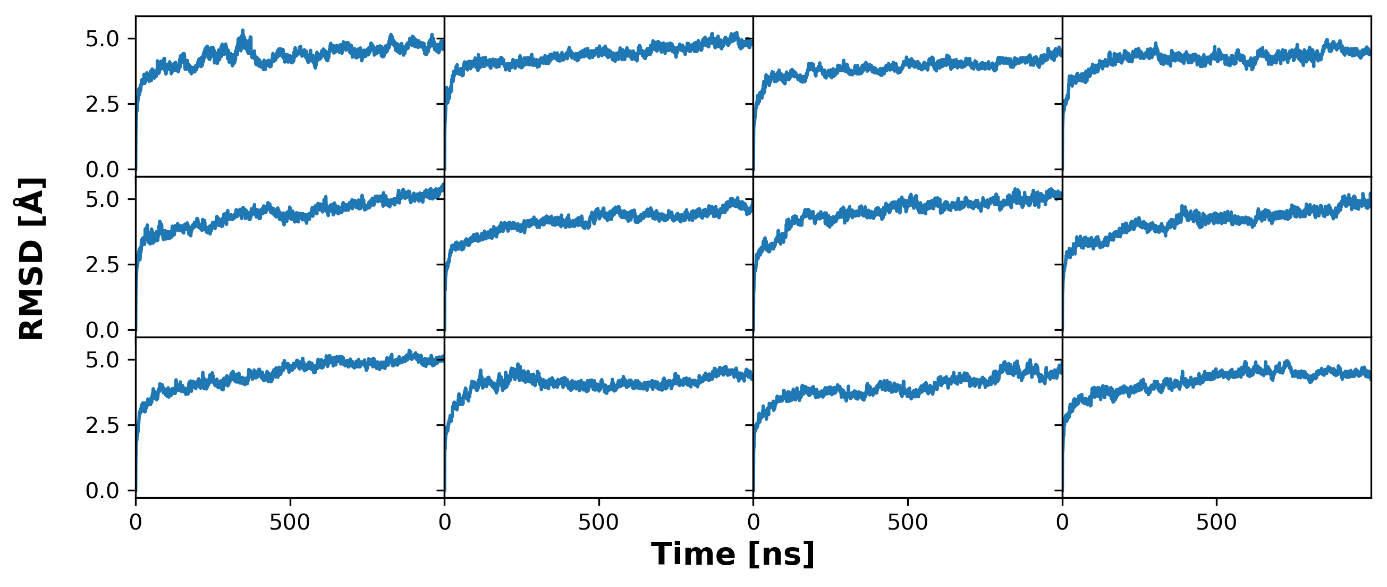

**Figure S4:** Backbone (C, CA, N) RMSD of 12 replicas of 1 μs long MD simulations of MB327 bound to the human nAChR compared to the first frame of each simulation.

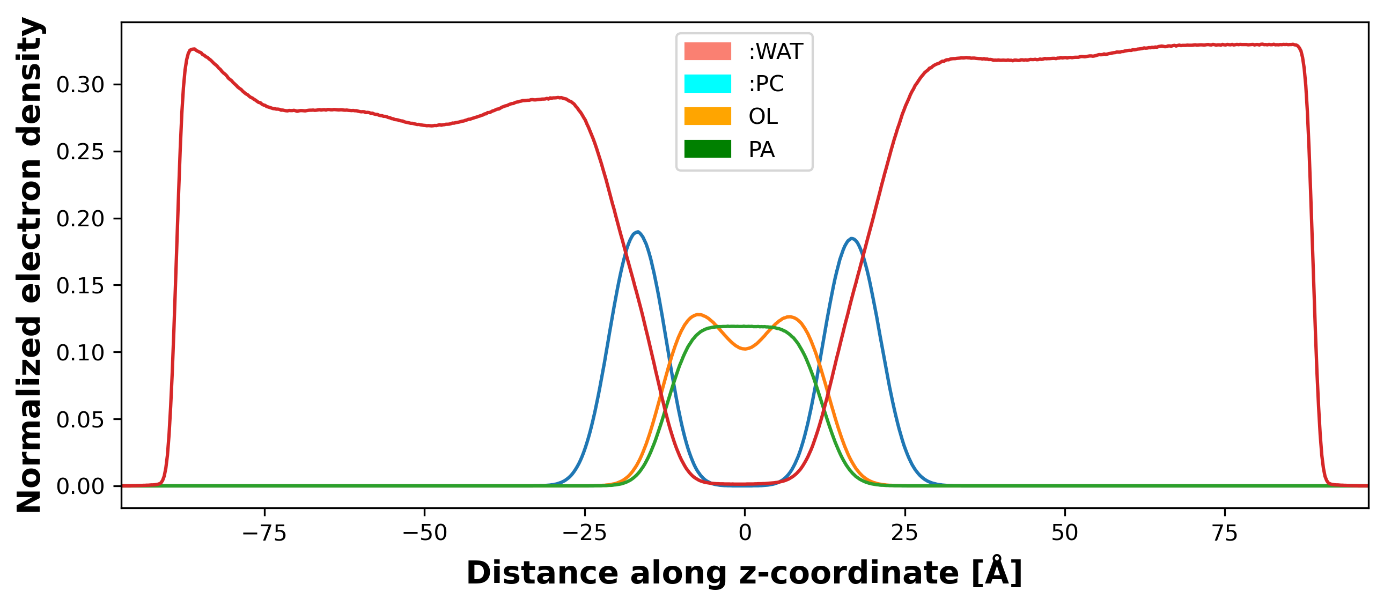

**Figure S5:** Normalized electron density of water and membrane components averaged over all replicas during MD simulations.

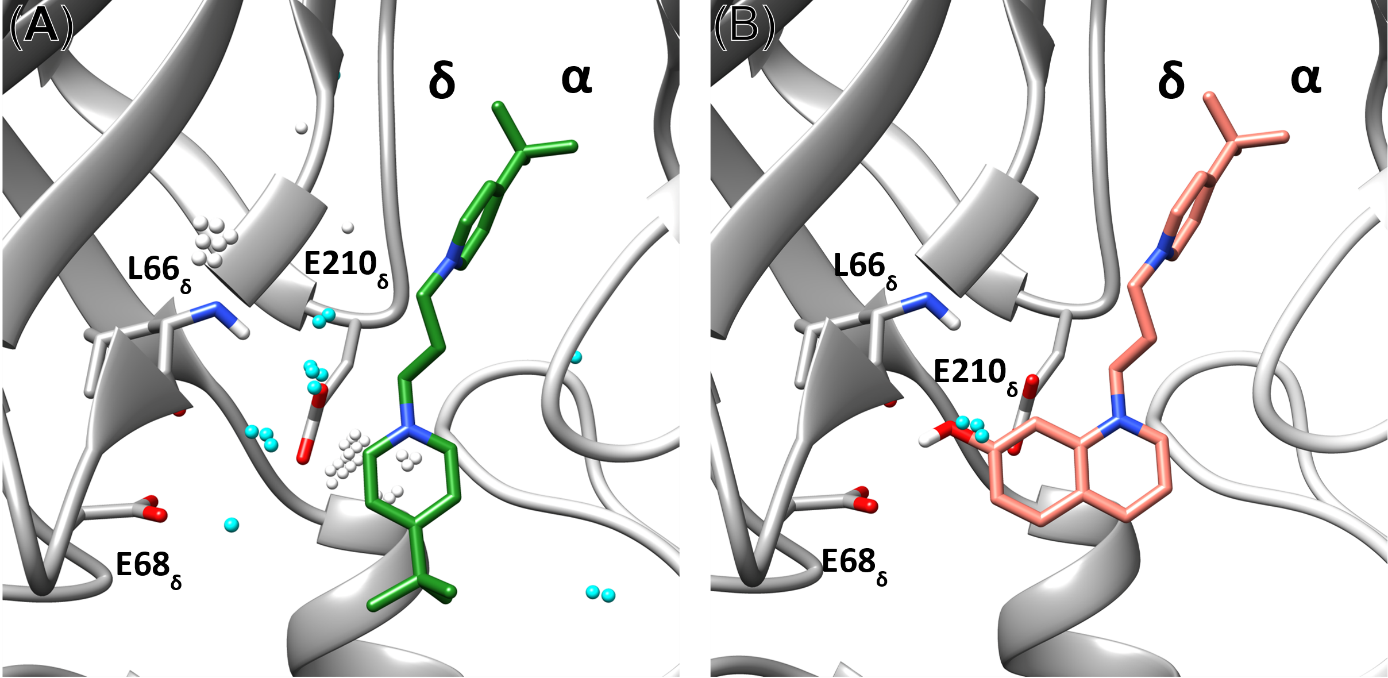

**Figure S6:** MB327 analog based on substituting water molecules in the allosteric MB327-PAM-1 binding pocket between the α- and δ-subunit. **A)** MB327 binding in between the α- and δ-subunit. Water clusters as identified by GIST are shown as spheres; clusters within 5 Å of MB327 are colored cyan. **B)** Proposed binding mode of PTMD90-0012 (salmon) in between the α- and δ-subunit. Modification of MB327 to PTMD90-0012 (salmon) leads to a substitution of a water cluster located in proximity to L66_δ_, E68_δ_, and E210_δ_. For L66 _δ_, the backbone atoms are shown in addition to the side chain.

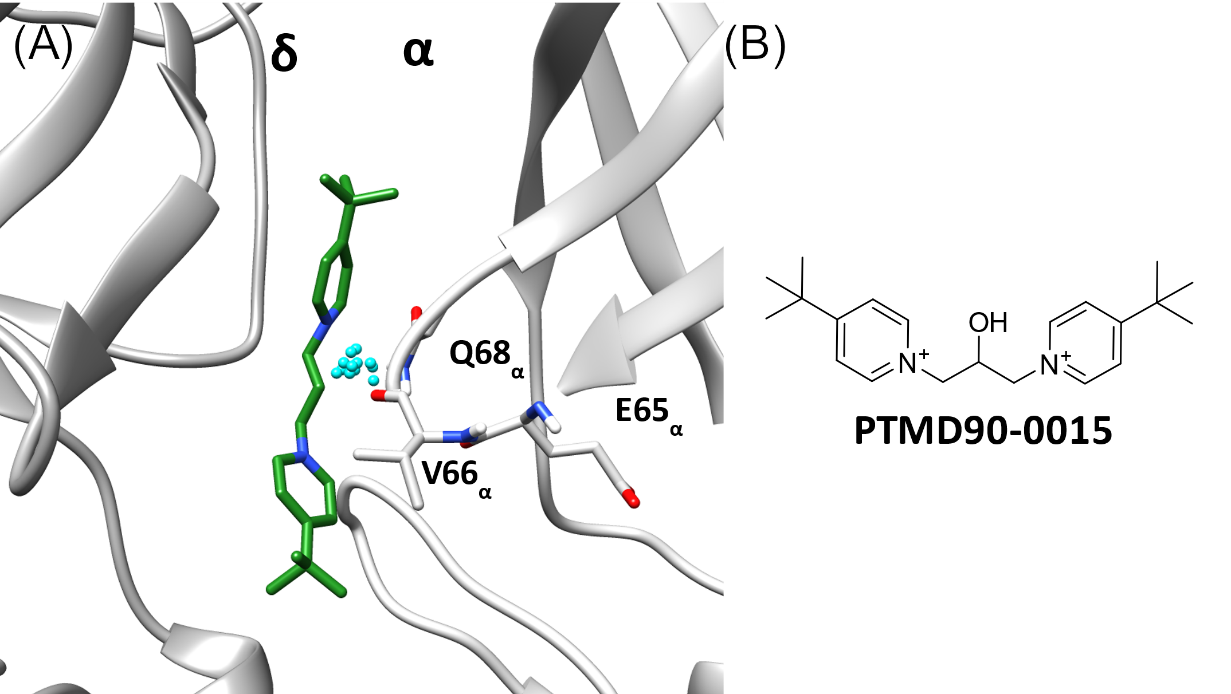

**Figure S7:** Initial water cluster leading to the design of PTMD90-0015. **A)** During the first few replicas (5 out of 12) of MD simulations of MB327 in MB327-PAM-1, an energetically unfavorable cluster of water molecules was identified in between the backbone atoms of E65_α_, V66_α_, and Q68_α_ close to the C3-linker of MB327 in a representative binding mode during MD simulations (k-means clustering based on MB327 atoms ligand between the α- and δ-subunit). Based on these preliminary results, **B)** PTMD90-0015 was designed. This water cluster was later only observed in between the δ- and α-subunit. However, there, no docked or representative structure from MD simulations was in the range to form interactions with the protein after modification of MB327 to PTMD90-0015.
